# Prediction of Motor Imagery Tasks from Multi-Channel EEG Data for Brain-Computer Interface Applications

**DOI:** 10.1101/2020.04.08.032201

**Authors:** Md. Ochiuddin Miah, Md. Mahfuzur Rahman, Rafsanjani Muhammod, Dewan Md. Farid

## Abstract

The classification of **m**otor **i**magery **e**l**e**ctroencephalo**g**ram (MI-EEG) is a pivotal part of the biosignal classification in the **b**rain-**c**omputer **i**nterface (BCI) applications. Currently, this bio-engineering based technology is being employed by researchers in various fields to develop cutting edge applications. The classification of real-time MI-EEG signal is the core computing and challenging task in these applications. It is well-known that the existing classification methods are not so accurate due to the high dimensionality and dynamic behaviors of the real-time EEG data. To improve the classification performance of real-time BCI applications, this paper presents a clustering-based ensemble technique and a developed brain game that distinguishes different human thoughts. At first, we have gathered the brain signals, extracted and selected informative features from these signals to generate training and testing sets. After that, we have constructed several classifiers using Artificial Neural Network (ANN), Support Vector Machine (SVM), naïve Bayes, Decision Tree (DT), Random Forest, Bagging, AdaBoost and compared the performance of these existing approaches with suggested clustering-based ensemble technique. On average, the proposed ensemble technique improved the classification accuracy of roughly 5 to 15% compared to the existing methods. Finally, we have developed the targeted brain game employing our suggested ensemble technique. In this game, real-time EEG signal classification and prediction tabulation through animated ball are controlled via threads. By playing this game, users can control the movements of the balls via the brain signals of motor imagery movements without using any traditional input devices. All relevant codes are available via open repository at: https://github.com/mrzResearchArena/MI-EEG.

## 1. Introduction

In this era, brain engineering is an emerging field of science and technology. It includes the apprehension and accomplishment of various fields like physics, chemistry, biology, computer science, and mathematics to base its application [1]. On top of that, it incorporates the concepts and methods of varied fields like biological science, clinical medicine as well as engineering [2]. Brain-Computer Interface (BCI) is a branch of brain engineering that targets to solve practical problems of the life sciences. BCI and Human Machine Interface (HMI) are the modern technology that is used as a tool to establish communication between users and machines [3]. BCI technology incorporates neurophysiological activities as input signals and then these signals interpret into meaningful commands employing Machine Learning (ML) algorithms to execute actions by an external system [4, 5]. BCI technology brings about a huge number of opportunities in the medical field especially for the treatment and rehabilitation for disabled people. Around the world, millions of people are affected by various forms of disability and physical impairment caused by various reasons such as – born disability, old-age disability, early-age disability, accidental/unforeseen issues, serious health problems, and more [6]. For disabled people, it is extremely difficult or, in some cases, impossible to do day to day chores without the assistance of any caregiver. A variety of equipment and devices invented in this field for rehabilitating and aiding disabled people to cope with their physical deficiencies [7]. This technology allows disabled people to command and control external devices such as – computers, wheelchairs, robots, etc., by utilizing their thoughts. Notwithstanding, the horizon of this field has been broadened to a wide range of non-medical applications like gaming, entertainment, military, and meditation training [8, 9]. The knowledge of BCI can also be applied in smart environment systems like smart houses or smart workplaces. This technology has opened a new window of opportunities for the gaming and entertainment industry and different types of games already being developed using this technology to relief from stress [10, 11].

Fundamentally, BCI technology can be employed in three ways based on the process of signal acquisition from brain [12]. One way is to place wires inside the grey matter of the brain, this is called invasive BCI [13, 3]. Though invasive BCI begets a better input signal, it is highly sophisticated and extremely sensitive to implement [12, 14]. That is why it necessitates very proficient people to manipulate with. Another way is to place electrodes on the surface of the scalp to measure activities from a huge group of neurons. This method is called non-invasive BCI [15, 16]. In comparison with invasive BCI, it does not deliver better input signals as good as [17]. But it is less sophisticated and easy to accomplish with. The third type of BCI is called partially invasive BCI. In this technique, wires are placed inside the brain but above the grey matter of it [12, 17].

Concerning motor imagery EEG signal classification, the proficiency of many single and ensemble classifiers had previously been evaluated. Our prior studies had assessed the performance of existing single and ensemble classifiers in real-time BCI applications [17, 13, 18]. It is well-known that the existing classifiers are not so accurate due to the high dimensionality and dynamic behaviors of the real-time EEG data [19, 20]. Sometimes, the signals are biased with artifacts and noise due to the low conductivity of the electrodes with the scalp [1]. The objective of this paper is to extend our prior works and ameliorates the classification performance by handling multiple electrodes/ neurons data at the same time. The proposed clustering-based ensemble technique clustered the dataset based on the position of the electrodes so that each cluster represents dissimilar information. It also selects the model dynamically based on the electrode locations to classify real-time EEG data. We have constructed several classifiers using Artificial Neural Network (ANN), Support Vector Machine (SVM), naïve Bayes, Decision Tree, Random Forest, Bagging, AdaBoost and compared the performance of these existing approaches with the suggested technique. We also developed a brain game to distinguish human thoughts in real-time. We employed the threading technique to control the signal classification and prediction tabulation via animated ball in real-time.

The remainder of this paper is organized as follows: Section 2 presents the related works. Section 3 illustrates brain measurement techniques, signal acquisitions, and datasets descriptions. Section 4 reveals supervised classification and the proposed clustering-based ensemble method. Section 5 provides experimental results and introduces a brain game that is controlled by brain signals of motor imagery tasks. Finally, Section 6 presents conclusions and future work.

## 2. Related Works

H. Raza et al. [1] presented an adaptive ensemble approach in EEG classification to handle non-stationarity in the motor imagery task; the proposed approach collected MI correlated brain responses and extracted spatial pattern structures from it, and new classifiers are added over time with an existing ensemble classifier to justify the deviations in streaming data input. Furthermore, BCI-related EEG datasets are employed to evaluate the proposed approach with single-classifier and it ameliorated the BCI performance significantly. S. Sreeja et al. [14] proposed a sparse based classification method to classify MI related tasks from EEG data. For enhancing the computational time, they only used wavelet energy without any pre-processing as a feature to classify MI data. In their experiments, the proposed sparsity approach outperformed existing classifiers in a minor computation time.

M. Li et al. [2] proposed a novel decoding approach employing Overlapping Averaging (OA) to interpret MI-EEG data. To overwhelm the constraint of the general Region of Interest (ROI) based decrypting approach, they used Weighted Minimum Norm Estimate (WMNE). They experimented on a public dataset employing 10-fold cross-validation and achieved higher decrypting accuracy of 81.32% than existing approaches. S. Sun et al. [19] researched to methodically assess the performance of popular ensemble methods: bagging, boosting and random subspace to classify MI related tasks from EEG signals. They used the base classifiers of SVM, DT, and k-Nearest Neighbors (k-NN) to advocate the possibilities of ensemble classification methods. Moreover, few significant inferences were produced to reveals the performance of ensemble methods for EEG classification in their study.

P. Pattnaik and J. Sarraf [21] attempted to apply left and right-hand movement classification using raw EEG signals. Before applying the movement classification, they had removed the artifacts in the obtained signals employing a low pass filtering technique. Moreover, they kept a strong focus on the application of BCI as well as the issues related to it. K. Mamun et al. [18] introduced a technique to procure frequency reliant neural synchronization as well as inter-hemispheric connectivity properties. They based their method on Granger causality as well as wavelet packet transform (WPT) approaches. Their approach was capable of decoding movement associated behaviors accurately as well as informatively, from the registered local field potentials (LFPs) activity. Movement recognition accuracy was 99.8%, and afterward laterality recognition was 81.5% on average. Finding of this study show that nominated optimum neural synchronization associated with inter-hemispheric connectivity and has the potential to give users control signal to supplement adaptive brain-machine interface (BMI).

J. Lu et al. [22] investigated the prospect of ameliorating performance in a Transcranial Doppler ultrasonography (TCD) based BCI. They used the structures and classifiers that are computationally suitable for online application by running an offline investigation of TCD recordings. From earlier offline TCD researches, pattern recognition and signal processing routine are used by them and received roughly 73% accuracy in their experiment. Later, methodical feature selection approaches and three classifiers such as: naïve Bayes, Linear Discriminant Analysis (LDA), and SVM were compared. Combining the SVM classifier with weighted sequential forward selection(WSFS), a topmost accuracy of 87.60±3.27% was acquired. M. M. Shanechi [23] examined the decoding algorithms made in the BMI study. It is possible to design a motor BMI as a closed-loop control system. They inspected current decoder designs that emphasize the unique properties of BMI. Moreover, a discussion was presented about the existing opportunities to formulate a control-theoretic framework to design BMI, as well as, assisting the development of more advanced BMI control algorithms.

R. M. Mehmood et al. [24] proposed a method that considerably enhances the rate of emotion recognition regarding the popularly implemented spectral power band routine. Features selected by this routine performed better than both univariate and multivariate attributes. The optimal attributes were later processed to classify emotion by applying KNN, SVM, LDA, naïve Bayes, Random Forest, and deep learning classifiers. M. A. Lebedev and M. A.L. Nicolelis[25] emphasized on some of the essential challenges encountered by BMI research. Moreover, they proposed a chain of milestones to convert up to date experimental advances into viable medical applications within the coming 10 - 20 years. The guideline they provided underscores the contemporary history of the BMI and puts a strong emphasis on the influential factors related to its growth.

## 3. EEG Signal Acquisition

### 3.1. Functional Areas of Brain

The human brain is the main organ of the human nervous system [18]. The central nervous system has consisted of both the brain and spinal cord. The human brain acts as the master of the whole body, as almost all of the body activities are controlled by it. It has the functionalities of receiving, processing and generating information for the well-being of human beings [26]. The sense organs send informations to the brain. The brain then integrates, coordinates and processes the given input informations to produce decisions and commands to the rest of the body [21]. Different brain parts are responsible for completing different tasks. Motor system is the part that both generates and controls the movements of the body [18]. The nerves do the job of transferring the motor system-generated movements from the brain to the motor neurons in the body. The action of the muscle is governed by the passing movements. Using the spinal cord, the corticospinal tract passes movements to the torso as well as to the limbs. The eyes, mouth, and face related movements are carried by the cranial nerve [25]. The motor cortex generates the movement of arms and legs. The motor cortex consists of three parts-primary motor cortex, premotor cortex, and supplementary motor area [22]. Locating on the frontal lobe of the brain, the primary motor cortex is one of the essential brain areas that are necessary for motor function. The primary motor cortex produces neural impulses and then these impulses control the execution of movement [13].

### 3.2. Brain Measurement Techniques

In BCI, different kinds of neurological modalities are applied to obtain neurological brain signals [18]. Electroencephalography (EEG) is one of the variety of ways that are applied for measuring brain activities. It is usually a non-invasive approach and assesses voltage oscillations ensuing from ionic current inside the brain neurons [12]. Another non-invasive method for measuring brain activities includes positron emission tomography (PET), magneto-encephalography (MEG), Transcranial Doppler ultrasonography (TCD) as well as functional magnetic resonance imaging (fMRI) [9, 27]. Some of the invasive electrophysiological methods are local field potentials (LFPs), electrocorticography (ECoG), as well as single-unit recording. EEG, ECoG, LFPs as well as single-neuron recordings are the only methods which are popular in use because of their relative simplicity and inexpensiveness, and they also come with high temporal resolution [18, 28]. On the other hand, fMRI, MEG, and PET are very costly and not much usable; which makes them unpopular to use [12]. In our experiment, we used Emotiv EPOC+ EEG neuroheadset because it is less sophisticated, inexpensive, and easy to use.

### 3.3. EEG Emotiv EPOC+ Neuroheadset

The Emotiv EPOC+ 14 channels is an EEG neuroheadset that ables to produce measurable electric potentials to assess brain activities [29]. It is equipped with 14 saline sensors (electrodes) which are put on the scalp of the brain following the international 10-20 system. In the 10-20 system, the real distance among two adjacent sensors can be either 10% or 20% [29]. The electrodes are situated in *F*_3_, *F*_4_, *FC*_5_, *FC*_6_, *F*_7_, *F*_8_, *AF*_3_, *AF*_4_, *T*_7_, *T*_8_, *O*_1_, *O*_2_, *P*_7_, *P*_8_ locations and two reference electrodes-Driven Right Lag (DRL) and Common Mode sense (CMS) are located at *P*_3_ and *P*_4_ locations [29, 13, 17]. The electrodes distribution of emotiv epoc+ employing the 10-20 system is shown in Fig. 1.

**Figure 1:**
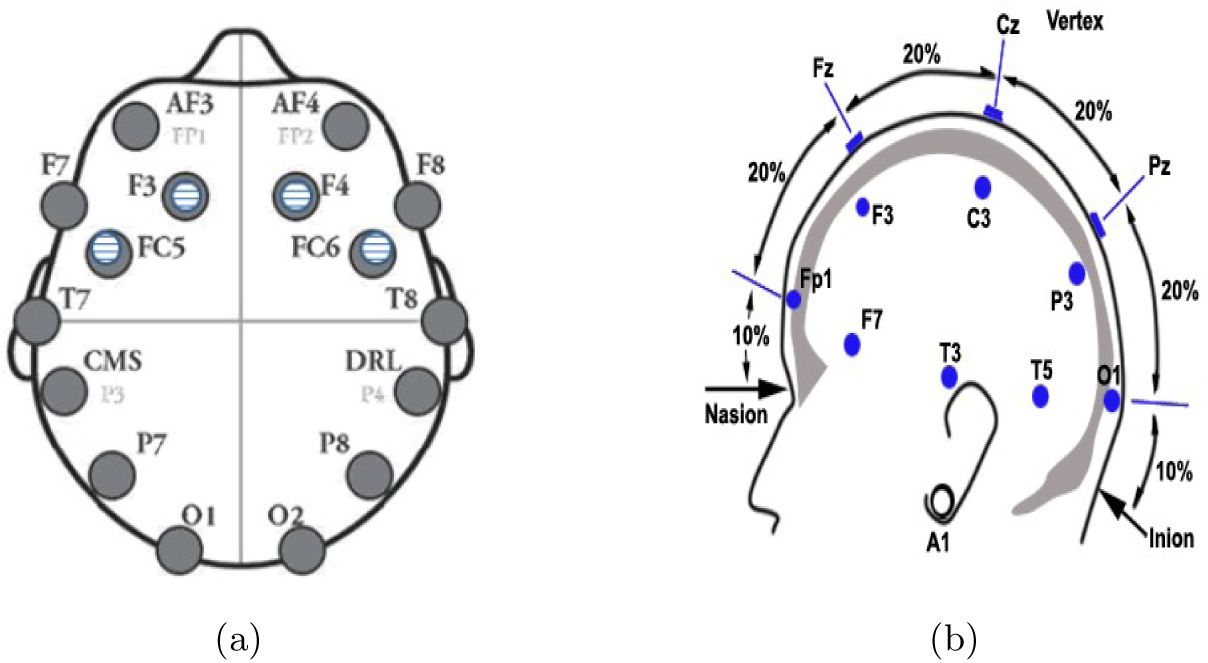
(a) Electrodes distribution of emotiv epoc+ neuroheadset (b) following the international 10-20 system. Source: emotiv systems.

### 3.4. EEG Brain Signal Recording

In this study, six healthy subjects (age 18±8) had participated and they had no neurological disorders. They were informed briefly about the experimental procedures and gave informed consent before the experiment. Firstly, we developed an application program via implementing Java-based technology as well as Emotiv SDK to obtain brain signals from Emotiv neuroheadset. Then, we acquired an average band power of theta, alpha, low beta, high beta, and gamma EEG neural frequency rhythms via the application program. EEG brainwaves, ranges and their association with different activities in the brain are described in section 3.5. We have mentioned earlier, the primary motor cortex produces neural impulses that control the execution of movements. In our MI hand movement experiment, we have selected *F*_3_, *FC*_5_, *FC*_6_ and *F*_4_ locations that are best fitted in the primary motor cortex area of the brain. We had employed Emotiv SDK API to acquire the average band power of these selected electrodes from the newest epoch with 0.5 seconds step size and 2 seconds window size. The average band power (BP) summarizes the overall power of the given frequency band and is calculated for the filtered band between 4 to 45 Hz by each electrode. We have recorded the MI-EEG brain signals for binary and ternary classes and the class values are steady, left and right-hand. During the experiment, participants wore the headset and connected it to the developed program computer through a wireless connection to the USB dongle. From each participant, we have taken two trials to obtain a training set for 30 seconds and one trial to acquire a testing set for 15 seconds. MI-EEG brain signal recording source code is available via open repository at: https://github.com/mrzResearchArena/MI-EEG/tree/master/Brain-Game/eeg_data_recording.

### 3.5. EEG Data Descriptions

The EEG measures different neural frequency rhythms that are associated with different regions, pathologies or brain states. Neural frequency is assessed by calculating the number of wave repeats within a second [12]. Table 1 illustrates the EEG brainwaves, ranges and their association with different activities in the brain [12, 3]. Most of the brain oscillations are connected with motor and sensory actions and associated with different brain functions. Brain oscillations are identified via electrodes then distributed into different frequency rhythms that are revealed in Fig. 2 [30, 28].

**Table 1:** EEG brainwaves, ranges and their association with different activities in the brain.

| Rhythms | Band (Hz) | Different band activities inside the brain |
| --- | --- | --- |
| Theta ($\theta$) | 4 to 8 | Sleeping, drowsiness, meditation, relaxed |
| Alpha ($\alpha$) | 8 to 12 | Calmness, learning, body integration, relaxed |
| Low_beta ($\beta$) | 12 to 18 | Active concentration, arousal, conscious thought |
| High_beta ($\beta$) | 18 to 25 | Anxiety, task-oriented, logical thinking |
| Gamma ($\gamma$) | 25 to 45 | Cognitive functioning and high processing tasks |

**Figure 2:**
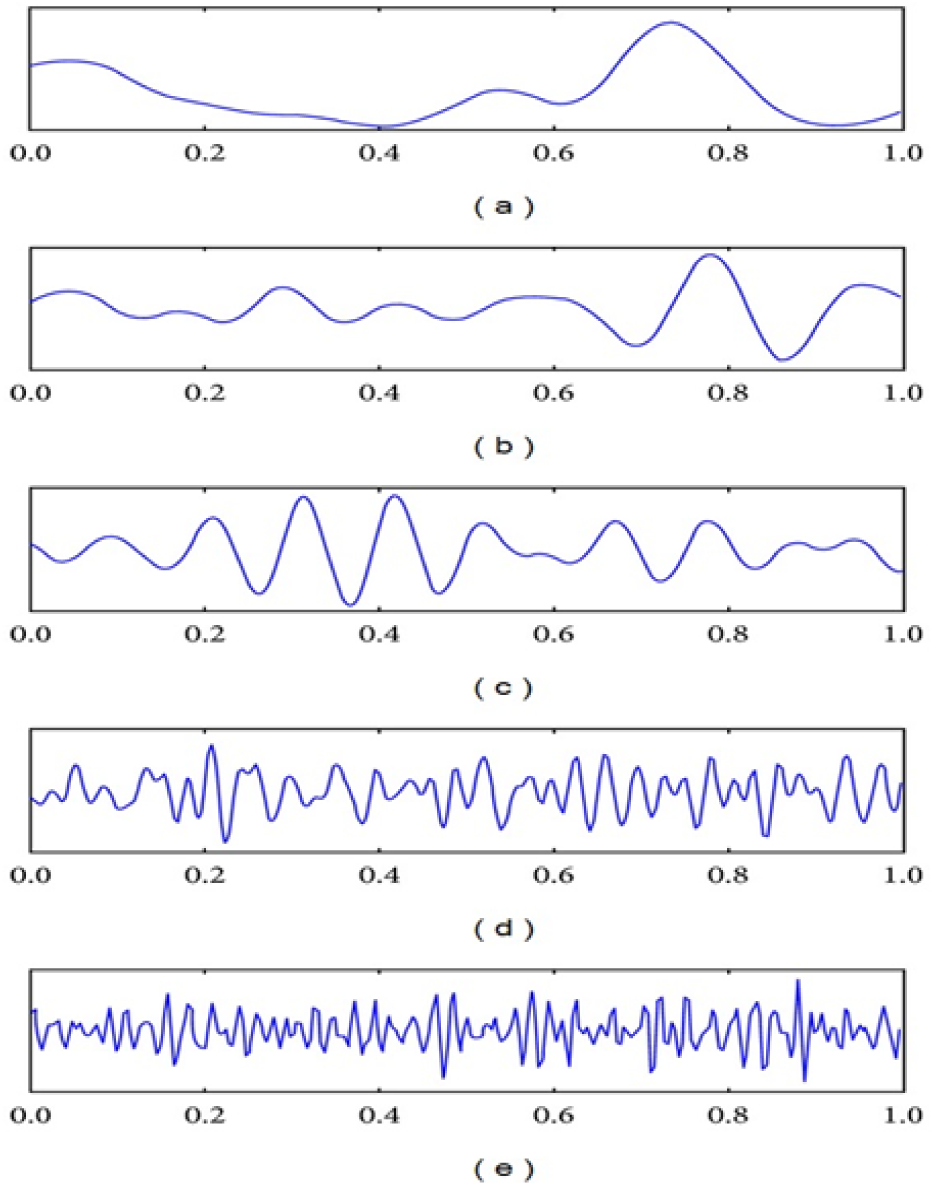
Different EEG neural frequency rhythms: (a) theta, (b) alpha, (c) low_beta, (d) high_beta, and (e) gamma

We have distributed the recorded MI-EEG dataset into binary-class and ternary-class sets based on the class values. We also occupied an EEG eye state dataset from the UCI Machine Learning repository (https://archive.ics.uci.edu/ml/datasets/EEG+Eye+State) to experience our proposed clustering-based ensemble method. This EEG dataset is collected via an emotiv epoc+ neuroheadset from the EEG measurement of two different eye states [31]. The comprehensive information of binary-class, ternary-class, and EEG eye state datasets are illustrated in Table 2.

**Table 2:**
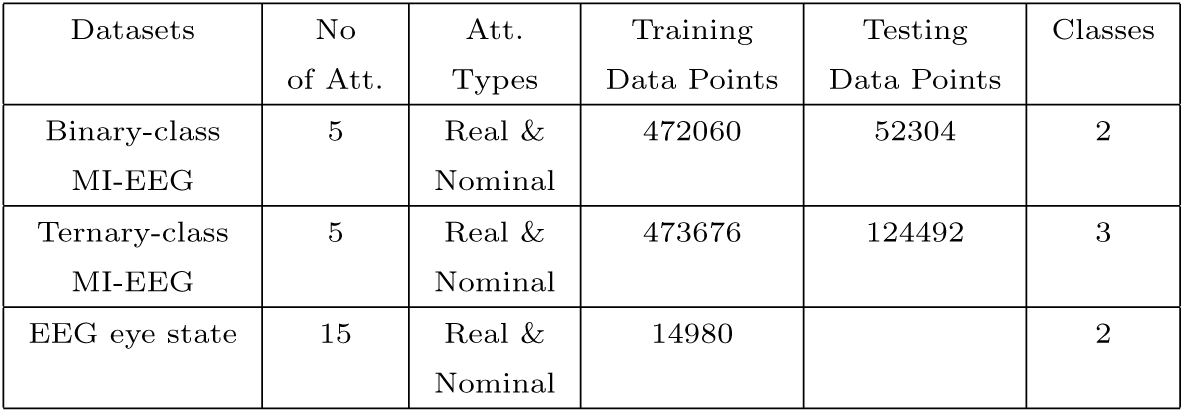
Hand movement MI-EEG and EEG eye state dataset discribtion.

| Datasets | No of Att. | Att. Types | Training Data Points | Testing Data Points | Classes |
| --- | --- | --- | --- | --- | --- |
| Binary-class MI-EEG | 5 | Real & Nominal | 472060 | 52304 | 2 |
| Ternary-class MI-EEG | 5 | Real & Nominal | 473676 | 124492 | 3 |
| EEG eye state | 15 | Real & Nominal | 14980 |  | 2 |

### 3.6. Visualization Brain Activities

The motor imagery hand movement deviations are found very high in the recorded MI-EEG responses as tabulated in Fig. 3. It represents that the movement and steady can be effortlessly found from the MI-EEG responses. It also highlighted that the movements of left and right-hand are challenging to categorize. We already mentioned that the primary motor cortex produces neural impulses to control the execution of movements. These movements are engendered from the related area and produced similar patterns that generate difficulty to classify patterns effortlessly.

**Figure 3:**
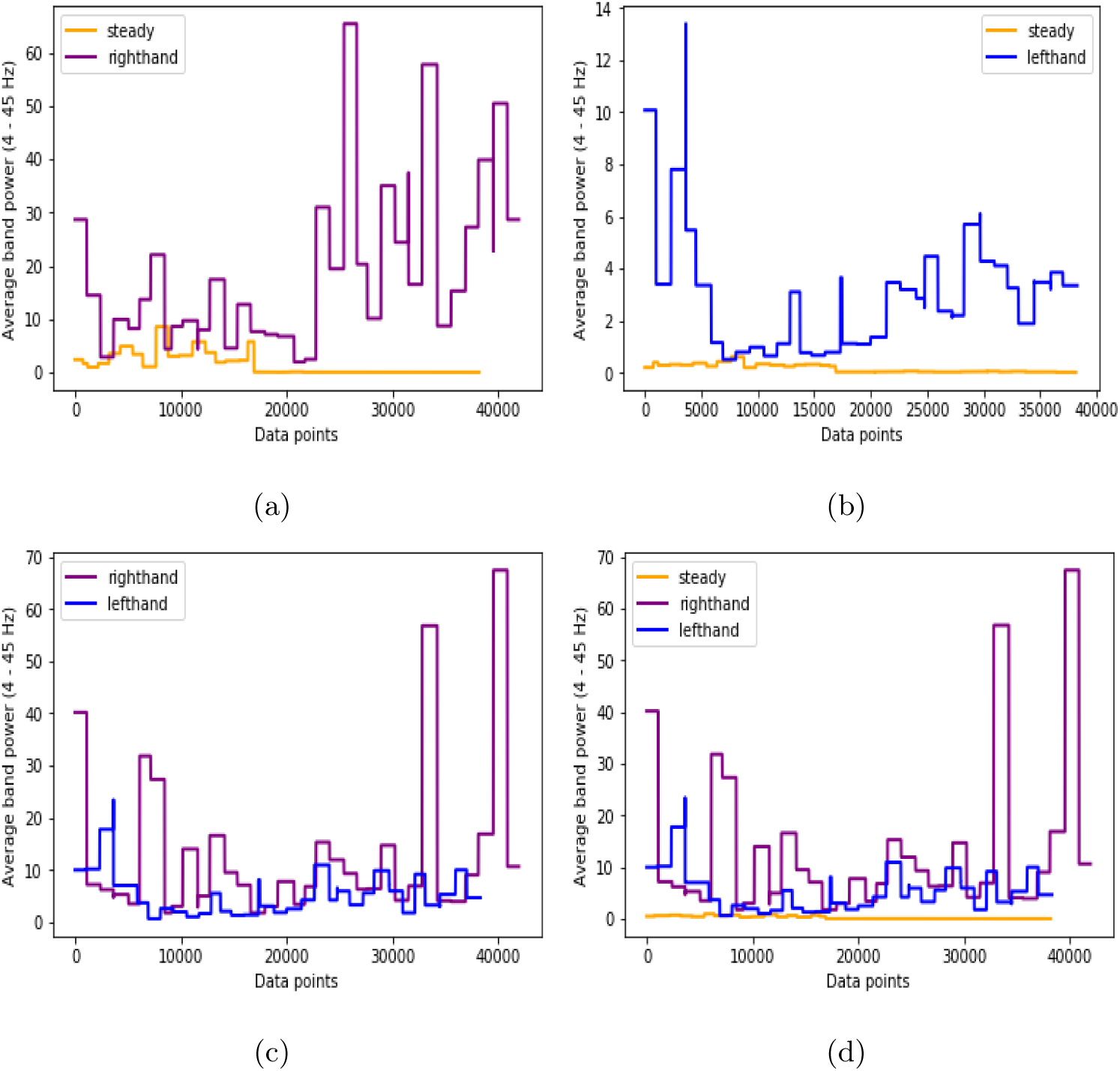
Dissimilar events inside the brain: (a) steady and right-hand movement; (b) steady and left-hand movement; (c) right and left-hand movement; (d) steady, right and left-hand movement.

## 4. Classification

In classification techniques, instance classification is accomplished via training and testing steps [32]. Training data points are used to build mining classifier, where all data points are labeled with class values. At the time of testing, data points are classified employing mining classifier [33]. In this technique, *N* number of instances reveal a dataset, *D* = {*x*_1_, …, *x*_*N*_} where each data point *x*_*n*_ ∈ *D* contains *F* features *A*_*f*_, *f* = 1, …, *F*. Here, 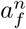 illustrates the value of attribute *A*_*f*_ of data point *x*_*n*_ and each one belongs to a class of *C* classes {*c*_1_, …, *c*_*l*_, …, *c*_*M*_}. In this section, we will briefly describe the proposed clustering-based ensemble model with four single classifiers and three ensemble learning methods.

### 4.1. Artificial Neural Network (ANN)

An artificial neural network (ANN) is a brain-inspired classification system based on the arrangement of biological neural networks [34]. It consists of input, output layers and sometimes single or multiple hidden layers exist to find complex patterns. To compute neuron input *I*, it uses bias weight *θ*, connected all neurons weight *W* and output *O* [34]. *After that, neuron input I* manipulates to calculate neuron output *O* employing different activation functions. Neuron input *I* and output *O* calculated by employing non-linear sigmoid function, are illustrated in Eq. 1 and 2 [24].

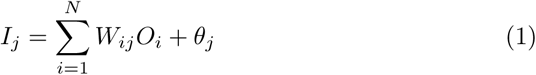

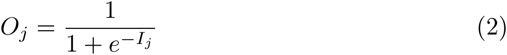

The network contains several connections, each connection computes the output of a neuron and then used by another neuron as input [24]. Weights are allocated to each connection to reveal relative importance. Initially, all the weights are assigned randomly. After that, each neuron learns from the set of training instances *D*, computes the error according to the desired output of *D* and adjusts the weights based on errors. The backpropagation algorithm is employed updating weights to ameliorate the network until achieving the pure output weight [34].

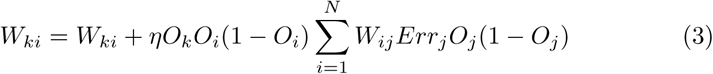

Eq. 3 is employed iteratively to adjust all the weights of the network. Here, the learning rate, *η* expresses the proportion of corrective steps to adjust the errors of the model in each iteration [34, 24].

### 4.2. Support Vector Machine (SVM)

A Support Vector Machine (SVM) is a supervised classifier that explores training instances and discovers hyperplanes, or support vectors to maximize the margin among the classes [35]. In two dimensional space, the support vectors split a plane into two chunks through a line where each cluster denotes individual classes [18]. A set of training data points, *D*= {*x*_1_, …, *x*_*N*_} having *N* number of data points with class values *C*= {1, −1} is employed to train the classifier. We can select two parallel support vectors with supreme probable distance to separate classes. The maximum distance between two hyperplanes is called the margin [18].

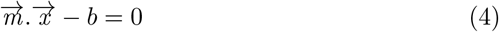

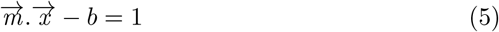

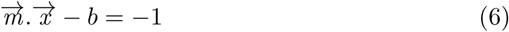

Any hyperplane can be inscribed as the Eq. 4 where 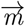 is a normal vector and parameter 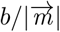 regulates the offset of the support vector from the origin [35]. Eq. 5 and 6 will be employed when the data points *x*_*n*_ are on or above the hyperplane and on or below the hyperplane respectively [23].

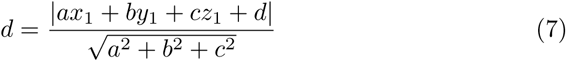

The distance, *d* among these two support vectors is 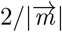. So, we have to minimize the 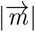 to maximize the *d*. The distance is calculated manipulating the distance among a point and plane equation that reveals in Eq. 7 [35, 18].

### 4.3. Naïve Bayes Classifier (NB)

Naïve Bayes classifier is a probabilistic machine learning classifier based on Bayes’ theorem. It engenders probability scanning the training instances only ones and can handle the missing attribute values easily by omitting its probabilities [32]. It takes a dataset, *D* with *F* number of attributes as input to build the classifier model. For testing a new instance, *x*_*new*_, the classifier will calculate the posterior probability and assign a class label with the highest probability.

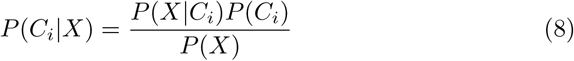

In Eq. 8, Bayes theorem states mathematically as *P*(*X*) is constant for all classes [32]. Here, *P* (*X*|*C*_*i*_) and *P*(*C*_*i*_) represents conditional and priori class probability. For instance, NB classifier predicts a new data point, *X* as class *C*_*i*_, if *P*(*X*|*C*_*i*_)*P* (*C*_*i*_) is greater than *P*(*X*|*C*_*j*_)*P*(*C*_*j*_) [33].

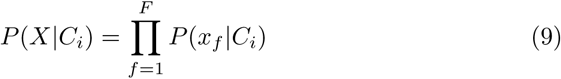

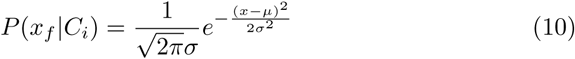

For data point *X, x*_*f*_ denotes to the value of feature *A*_*f*_. The probability *P* (*x*_*f*_ |*C*_*i*_) can be simply assessed from the training instances, if feature, *A*_*f*_ is categorical-valued. Otherwise, feature *A*_*f*_ is continuous-valued, then *A*_*f*_ is estimated through a Gaussian distribution with standard deviation *σ* and mean value *µ* shown in Eq. 10 [33, 32].

### 4.4. Classification and Regression Tree (CART)

The CART uses *Gini Index* that engenders binary classification tree to make decisions [32]. Firstly, it assesses the adulteration of dataset, *D* where probability *P*_*n*_ is estimated through |*c*_*l*_, *D*|*/*|*D*| reveals in Eq. 11 [33]. Here, the sum is calculated over *C* classes and each instance, *x*_*n*_ ∈ *D* belongs to a class *c*_*l*_.

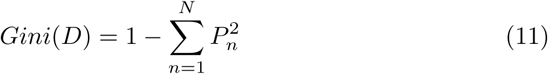

It splits the dataset, *D* considering binary split and the weighted sum of the adulteration of every resulting sub-data. For instance, the dataset, *D* splits into *D*_1_ and *D*_2_ considering the *Gini Index* of *D* which is calculated in given Eq. 12 [36].

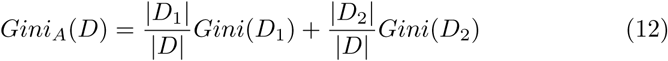

For each attribute, *A*_*f*_ considers each probable sub-data and nominated as the splitting attribute which ameliorate the reduction impurity, shown in Eq. 13 [36, 33].

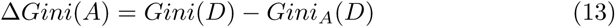

According to the attribute *Gini Index* impurity, it will split the dataset and create leaf nodes until all the splitting data belongs to an equivalent class. Algorithm 1 outlines the Classification and Regression Tree Algorithm [37].

#### Algorithm 1 CART Algorithm

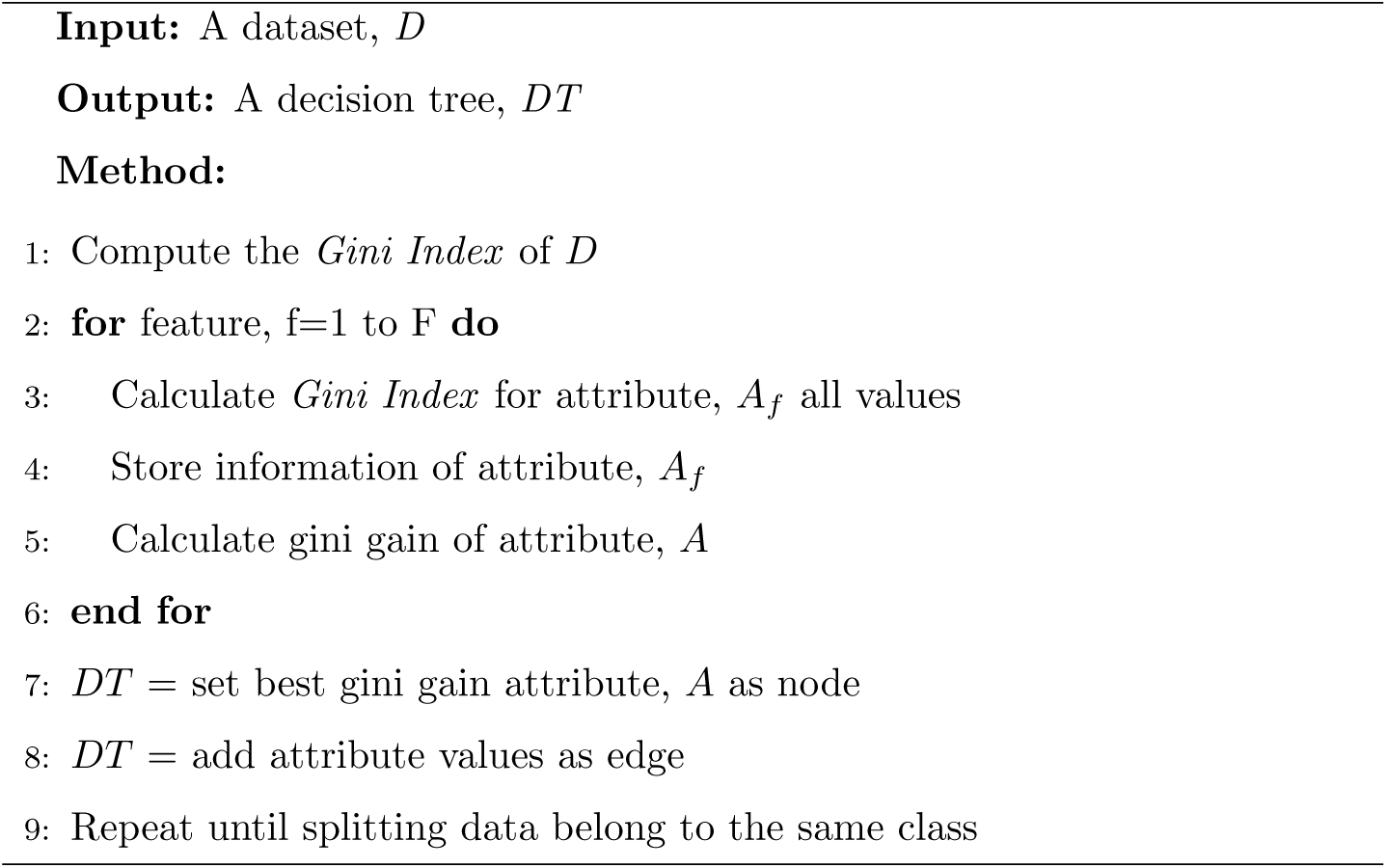

### 4.5. Random Forest

Random forest is an ensemble approach to classify a high volume of data with superior accuracy [38]. Initially, it splits a set of instances, *D* into numerous subsets *D*_1_, *D*_2_, …, *D*_*n*_ depending on the number of features *F*. Then, it constructs multiple decision trees based on the number of subsets and returns the most popular class among the trees as prediction [38, 19]. The learning scheme of random forest is tabulated in Algorithm 1. There is an association between the accuracy and number of engendered trees, the number of trees is directly proportional to accuracy. In machine learning, overfitting is one of the crucial problems and it may abate the classifier accuracy [38]. Random Forest classifier has overcome this difficulty. As it considers the vote of generated every decision trees, it will not overfit the model and gain greater accuracy than a single classifier model [38, 1].

### 4.6. Bagging

Bagging is also called Bootstrap Aggregation, which is an ensemble approach employed in statistical classification and regression to boost the performance of ML algorithms [1]. It combines the prediction of different equal-weighted models and classifies a new instance using the voting technique[19]. In the bagging technique, it requires a set of instances, *D* of size *N*, number of iterations, *I* to build the classifier model. It generates new datasets, {*D*_1_, *D*_2_, …, *D*_*n*_} by sampling the original dataset, *D* with replacement until the iterations number, *I*. Then, it uses each sub-datasets and learning scheme illustrated in Algorithm 1 to derive classifier models. To classify a new instance, *x*_*new*_, it combines the output of each model and considers the majority voting as prediction [19, 1, 39].

### 4.7. AdaBoost

Adaptive Boosting (AdaBoost) is a popular ML meta-algorithm, which combines a series of classifiers weighted votes to classify identified or unidentified instances [1]. It emphases on noisy data and builds a strong classifier by combining a set of weak classifiers [40]. Initially, it assigns an equal weight, 1*/d* to each training data point, *x*_*i*_ ∈ *D*. Then, it engenders a group of datasets, {*D*_1_, *D*_2_, …, *D*_*n*_} by sampling *D* with replacement based on instance weight until the iterations number, *I*. Each generated dataset, *D*_*i*_ derives a model, *M*_*i*_ and computes the error of the model by adding the weights of all instances in *D*_*i*_. Eq. 14 is shown the error calculation function of a model where *err*(*x*_*j*_) will be 1 when *x*_*j*_ will misclassified, otherwise 0 [41, 42].

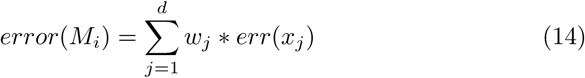

If the error of a model exceeds 0.5, we will regenerate *D*_*i*_ and derive a new *M*_*i*_. We will update the weight of an instance as if the weight of classified instances is abated and misclassified instances are enlarged [37]. To classify a new instance, it combines the votes and weights of each classifier.

### 4.8. Proposed Ensemble Method

The ensemble method is a technique to construct a powerful model by incorporating numerous classifiers. It considers several classifiers prediction to acquire state-of-the-art performance. In this section, we will discuss the proposed clustering-based ensemble technique that is used to classify motor imagery hand movement tasks. Initially, the proposed method takes the training EEG brain data *D*= {*x*_1_, …, *x*_*N*_} from *NE* neurons, or electrodes. Then, we have clustered the dataset, *D* into *NE* clusters based on the position of the electrodes. All clustered datasets *D*_1_, *D*_2_, …, *D*_*NE*_ represent diverse electrodes and labeled with previous class values. We also clustered the attributes of every sub dataset, *D*_*n*_, into *C* clusters. After getting sub dataset, we have construct *m* number of decision trees *D*_1_, .., *DT*_*m*_ with *m*^*th*^ cluster attributes employing decision tree induction algorithm (CART). The working procedures of the CART algorithm reveals comprehensively in section 4.4. We compute the error rate of *DT*_1_, *DT*_2_, …, *DT*_*m*_ on dataset, *D*_*n*_ and consider the minimum error rate as threshold, *T*. Then, we have considered the decision tree, *DT*_*m*_ with minimal error rate for ensemble trees *DT* ^∗^. Finally, we predict each real-time EEG data point, *x*_*new*_ employing *DT*^∗^ based on the position of the electrodes and deliberate the majority vote as prediction among the predictions of *DT*_*n*_ ∈ *DT*^∗^. The data points are coming from dissimilar electrodes and select the model dynamically based on the position of the electrodes. The process of classifying motor imagery tasks is illustrated in Fig.4 and the proposed clustering-based ensemble method is summarized in Algorithm 2. Space and time complexity of the proposed method depends on the number of features, dimensions of training data and size of the engendered tree.

**Figure 4:**
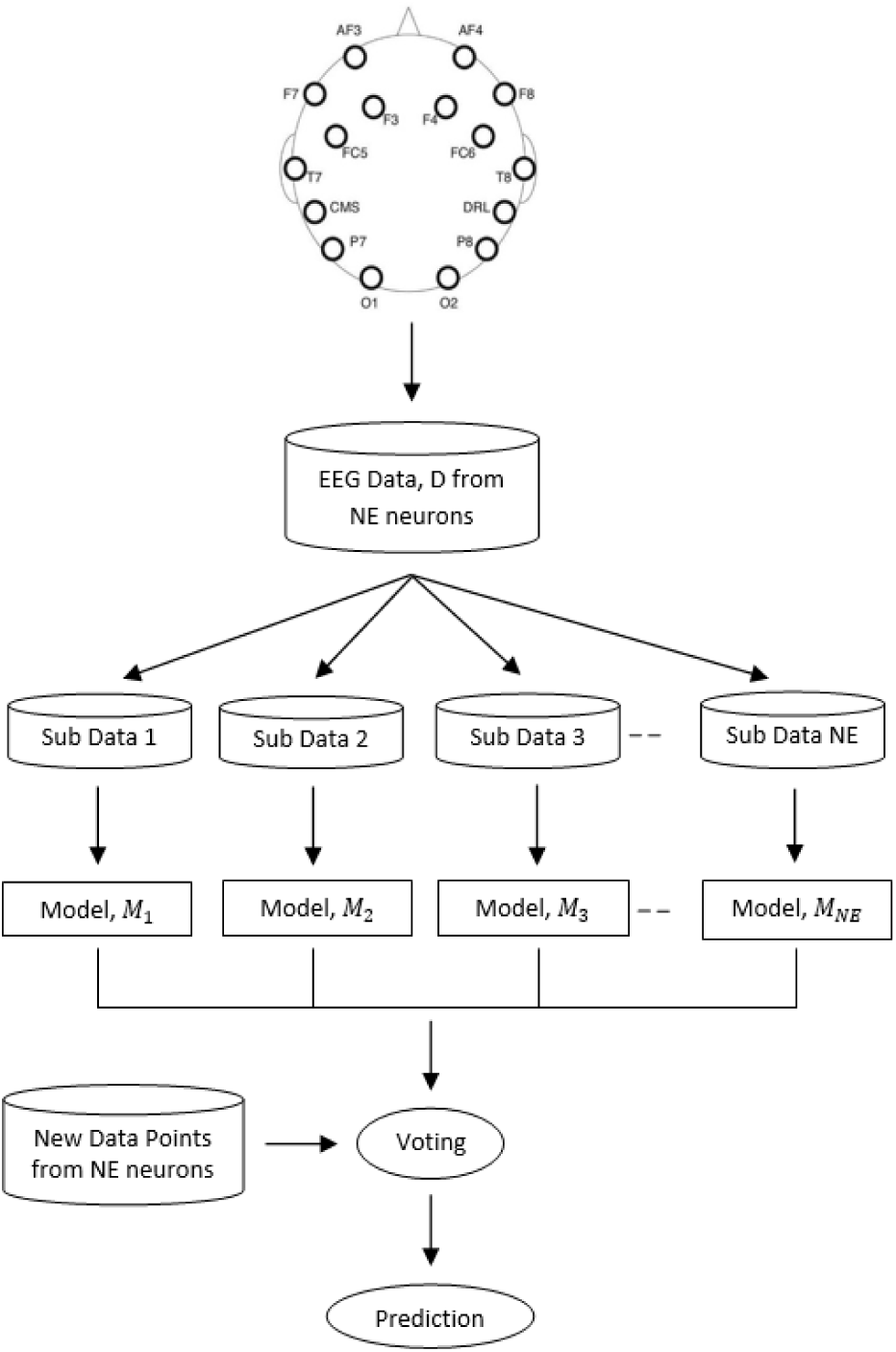
The process of classifying real-time motor imagery EEG data.

#### Algorithm 2 Proposed Clustering-based Ensemble Method

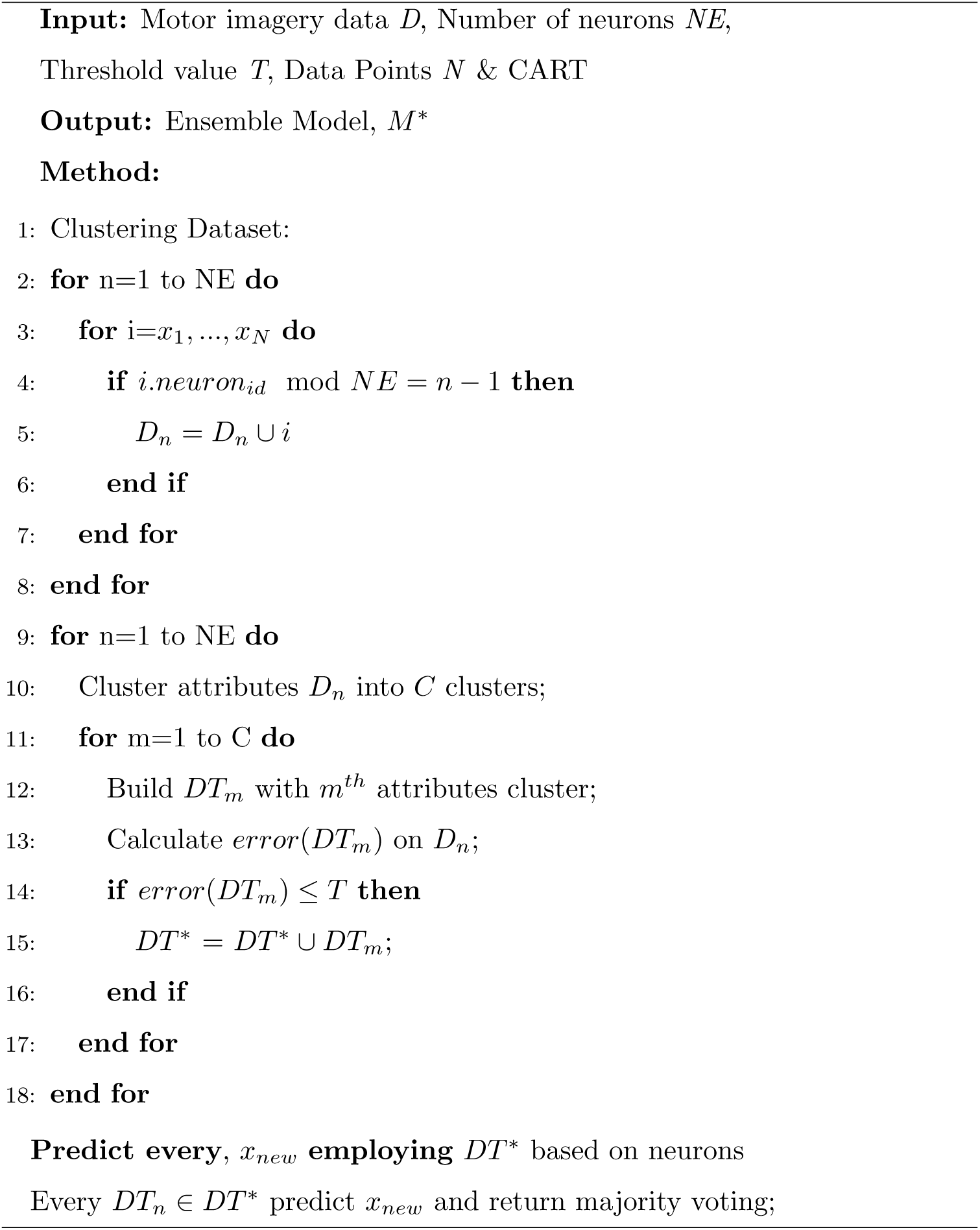

In BCI, real-time MI related EEG signal classification is a challenging task. Sometimes, the signals are biased with artifacts and noise due to the low conductivity of the electrodes with the scalp [1]. Different areas of the brain are responsible for the individual task and each electrode on the scalp provides dissimilar signals [19]. To ameliorate the performance of EEG signal classification in real-time is a demanding job because of the high dimensionality of the data and dynamic behavior of the electrodes [40]. If the training set is high dimensional, it will be challenging to build a good model employing single classifiers like ANN, SVM, naïve Bayes, and Decision Tree [19, 20]. Ensemble learning methodologies have been employed widely to grab these challenges. But existing ensemble methods generate sub-datasets by sampling the original dataset with replacement technique [1]. By applying this technique, the same instance can be repeated several times. The proposed clustering-based ensemble technique is to overcome these problems and classifies MI related EEG signals in real-time. It clustered the dataset based on the position of the electrodes so that each cluster represents dissimilar information. It also selects the model dynamically based on the electrode locations to classify real-time EEG data.

## 5. Experiments

In this section, we will describe the experimental environments, results of the proposed clustering-based ensemble method and present the developed brain game that is controlled by real-time motor imagery hand movements data.

### 5.1. Experimental Setup

In this study, the experiments were conducted via a device with an Intel Core-i5 (2.60 GHz) processor, and 8 GB of RAM. We implement the proposed method in Python programming language (version 3.7) and used the scikit-learn (version 0.21.2) as a machine learning library. We have tested the performance of the proposed method with some popular machine learning algorithms using classification accuracy, precision, recall, and F-score. The accuracy is measured by Eq. 15 where *assess*(*x*_*i*_) = 1 when *x*_*i*_ is correctly classified or *assess*(*x*_*i*_) = 0 when *x*_*i*_ is misclassified [32, 36]. The weighted average values of precision, recall, and F-score are considered and the calculations are shown in Eq. 16 to 18 [36, 37]. We also represent decision boundaries using the area under the ROC curve and AUC score. In the ROC curve, true positive rate (TPR) is plotted against false positive rate (FPR) and the calculations are defined in Eq. 19 and 20 [13]. The lowest threshold is considered through a line, *y* = *x* in au-ROC curve where correctly classified data points represent 1 and misclassified instances reveal as 0 [38].

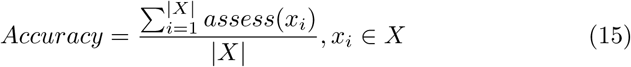

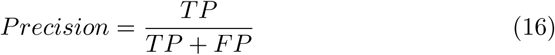

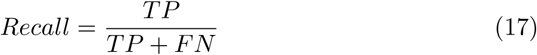

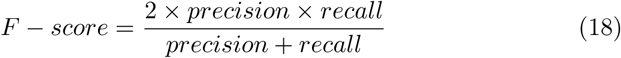

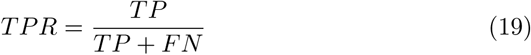

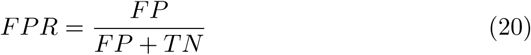

Here TP, TN, FP, and FN indicate the number of positive samples correctly classified, negative samples correctly classified, negative samples incorrectly classified and positive samples incorrectly classified respectively [36, 37].

### 5.2. Results

Firstly, we evaluated the performances of the proposed method against existing ANN, SVM, naïve Bayes, Decision Tree, Random Forest, Bagging and AdaBoost Classifiers on training sets of motor imagery EEG datasets. We have tested the classifiers model using Python Scikit-learn machine learning libraries supplying each of the testing EEG data.

The results in Table 3 point out that the proposed clustering-based ensemble algorithm outperforms the existing classifiers on a binary-class motor imagery EEG dataset. The proposed method performed superior and reached 99% accuracy on average for binary-class dataset. In this dataset, single classifiers are failed to achieve more than 85% accuracy where existing ensemble approaches can achieve 91% accuracy. The decision boundaries of the proposed method with some existing classifiers for this dataset is shown in Fig. 6.

**Table 3:**
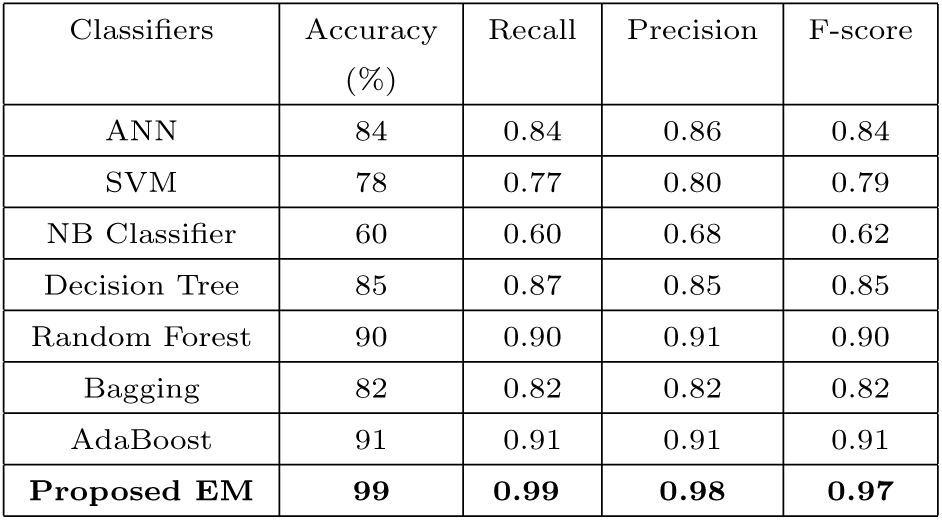
Performance comparison of the proposed clustering-based ensemble model with some popular algorithms on binary-class EEG dataset.

**Figure 5:**
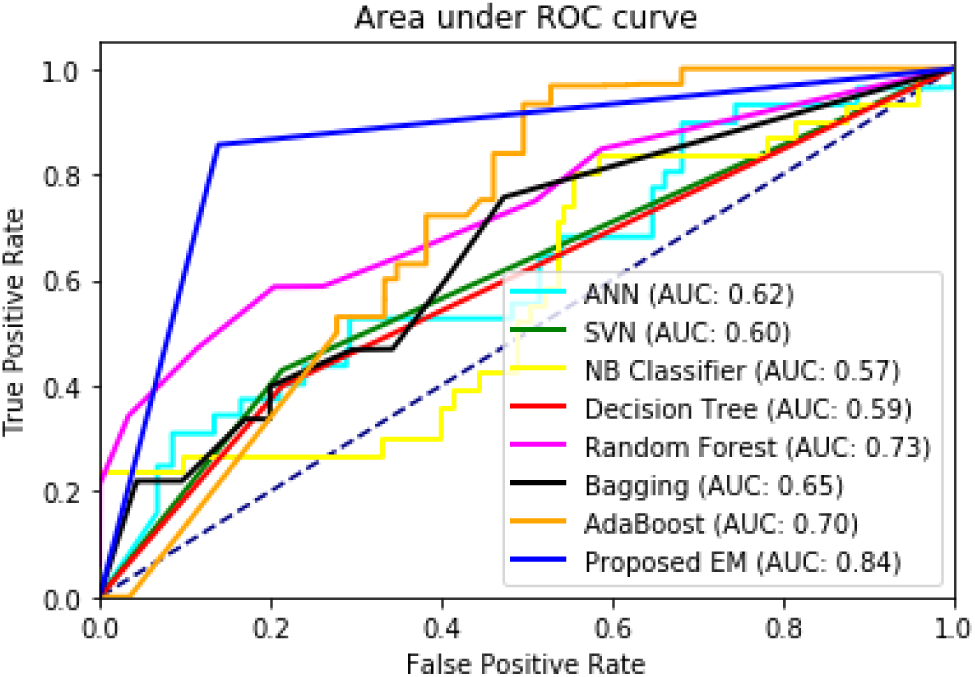
ROC and AUC analysis of the proposed method with some existing algorithms on ternary-class motor imagery EEG dataset.

**Figure 6:**
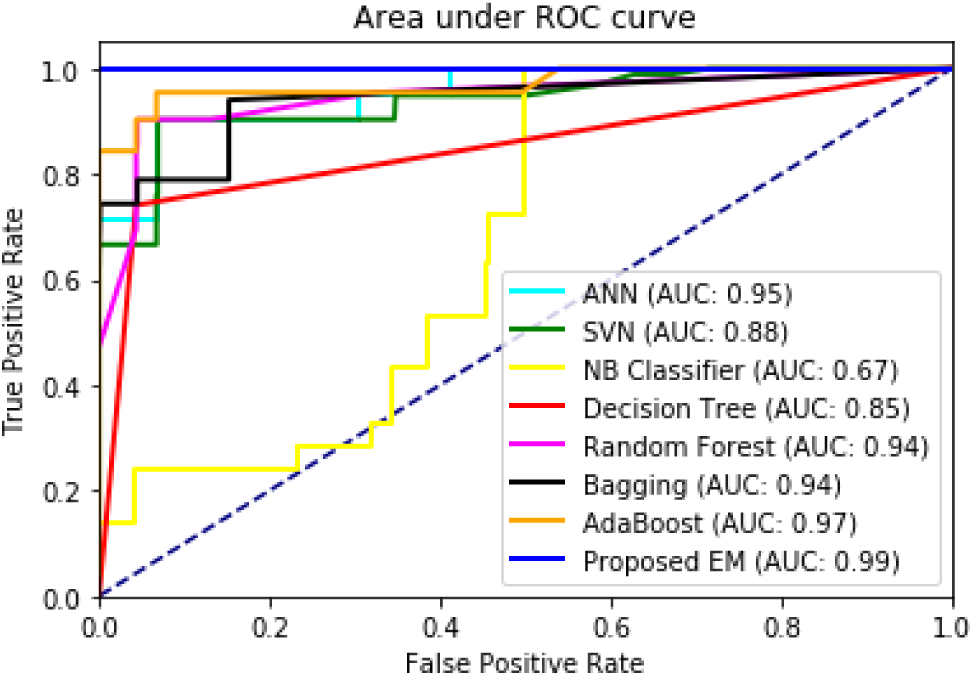
ROC and AUC analysis of the proposed procedure with some popular algorithms on binary-class MI-EEG dataset.

Moreover, in Table 4, the proposed clustering-based ensemble classifier also outperforms some popular machine learning algorithms on ternary-class dataset. In the ternary-class dataset, left and right-hand movements classification were challenging because both tasks are engendered from the motor cortex and samples are associated. This time our proposed method achieved better than the existing four single classifiers and three ensemble methods. Fig. 5 illustrates the decision boundaries of the classifiers via ROC analysis and AUC scores for the ternary-class dataset.

**Table 4:** Performance comparison of the proposed clustering-based ensemble model with some popular algorithms on the ternary-class EEG dataset.

| Classifiers | Accuracy (%) | Recall | Precision | F-score |
| --- | --- | --- | --- | --- |
| ANN | 53 | 0.53 | 0.55 | 0.54 |
| SVM | 57 | 0.55 | 0.57 | 0.56 |
| NB Classifier | 60 | 0.60 | 0.64 | 0.59 |
| Decision Tree | 54 | 0.54 | 0.55 | 0.54 |
| Random Forest | 62 | 0.62 | 0.64 | 0.61 |
| Bagging | 57 | 0.57 | 0.59 | 0.57 |
| AdaBoost | 61 | 0.62 | 0.63 | 0.59 |
| **Proposed EM** | **79** | **0.79** | **0.80** | **0.79** |

Besides, we used the EEG eye state dataset to experience our proposed clustering-based ensemble method. For this dataset, we have also significant upgrading of our proposed algorithm and achieved 90% accuracy on average.

Decision Tree performs average and reached 83% accuracy where existing ensemble approaches achieved 89% accuracy. The comparison of accuracy, precision, recall and F-score analysis using 10-fold cross-validation for EEG eye state dataset are tabulated in Table 5. Fig. 7 reveals the ROC analysis and AUC scores of the proposed method for the EEG eye state dataset.

**Table 5:** Performance comparison of proposed clustering-based ensemble model with some popular algorithms on EEG eye state dataset.

| Classifiers | Accuracy (%) | Recall | Precision | F-score |
| --- | --- | --- | --- | --- |
| ANN | 44 | 0.43 | 0.44 | 0.43 |
| SVM | 64 | 0.63 | 0.63 | 0.63 |
| NB Classifier | 54 | 0.55 | 0.55 | 0.42 |
| Decision Tree | 83 | 0.84 | 0.84 | 0.84 |
| Random Forest | 88 | 0.89 | 0.90 | 0.90 |
| Bagging | 89 | 0.90 | 0.90 | 0.90 |
| AdaBoost | 75 | 0.75 | 0.75 | 0.75 |
| **Proposed EM** | **90** | **0.90** | **0.91** | **0.90** |

**Figure 7:**
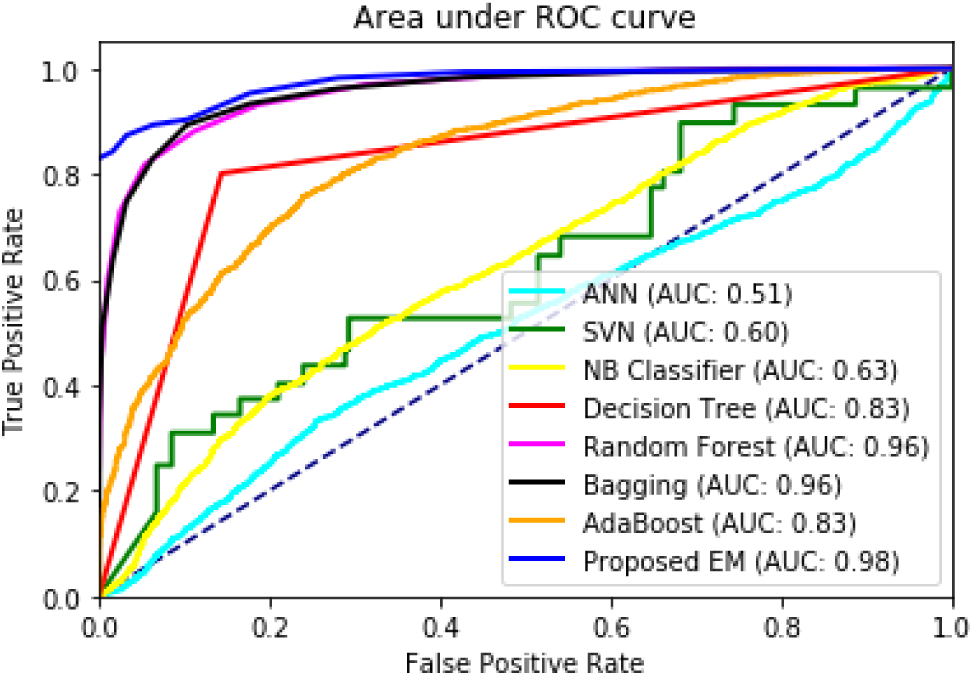
ROC and AUC analysis of the proposed ensemble technique with some existing methods on EEG eye state dataset.

Overall, algorithm 2 is performed better than several traditional data mining algorithms as well as achieved high accuracy and AUC score on average on three EEG datasets. The proposed method delivered an enhancement of roughly 10 to 20% accuracy parallel to single classifiers. It also ameliorated the classification accuracy of approximately 5 to 15% compared to existing ensemble approaches. Although the proposed clustering-based ensemble method outperforms other existing classifiers, there are limitations to be considered. Sometimes, the input signal is influenced by artifacts and noise due to the short conductivity of the neurons with the scalp. For these reasons, the classifier failed to correctly classify data points in real-time and the performance of the classifier is abated.

### 5.3. Developing Brain Game

We used our proposed clustering-based ensemble model and Java-Swing technology to develop our targeted application (game) system. We also used the emotiv community SDK in our Java program to acquire live brain signals. This game takes average band power of different frequency bands from *F*_3_, *FC*_5_, *FC*_6_ and *F*_4_ electrodes as input commands. Then these commands are classified by the proposed model based on the position of the electrodes and provide some actions as prediction. The input commands are coming from *F*_3_, *FC*_5_, *FC*_6_, *F*_4_ electrodes and select the model dynamically based on the position of the electrodes. We have exposed some actions via animated balls according to the predictions and percentage of different classified class. The real brain data classification through the developed model and prediction tabulation via animated balls in real-time is controlled via threads. The final prediction is taken from the dynamic number of instances votes to control the game more precisely. We have developed two versions of this game and they can identify binary as well as ternary actions of motor imagery task without any necessity of conventional input devices. It also increased the classification accuracy of real-time EEG signals of motor imagery tasks. The animated balls in Fig. 8 are presenting the movement and steady correspondingly. This application system can be used for rehabilitation as well as upgrading of user well-being. The user can exercise his concentration to recover from attention deficiency and boost his attentions via playing this brain game. It can also be used for gaming and entertainment purposes. The source code of brain game is available via open repository at: https://github.com/mrzResearchArena/MI-EEG/tree/master/Brain-Game.

**Figure 8:**
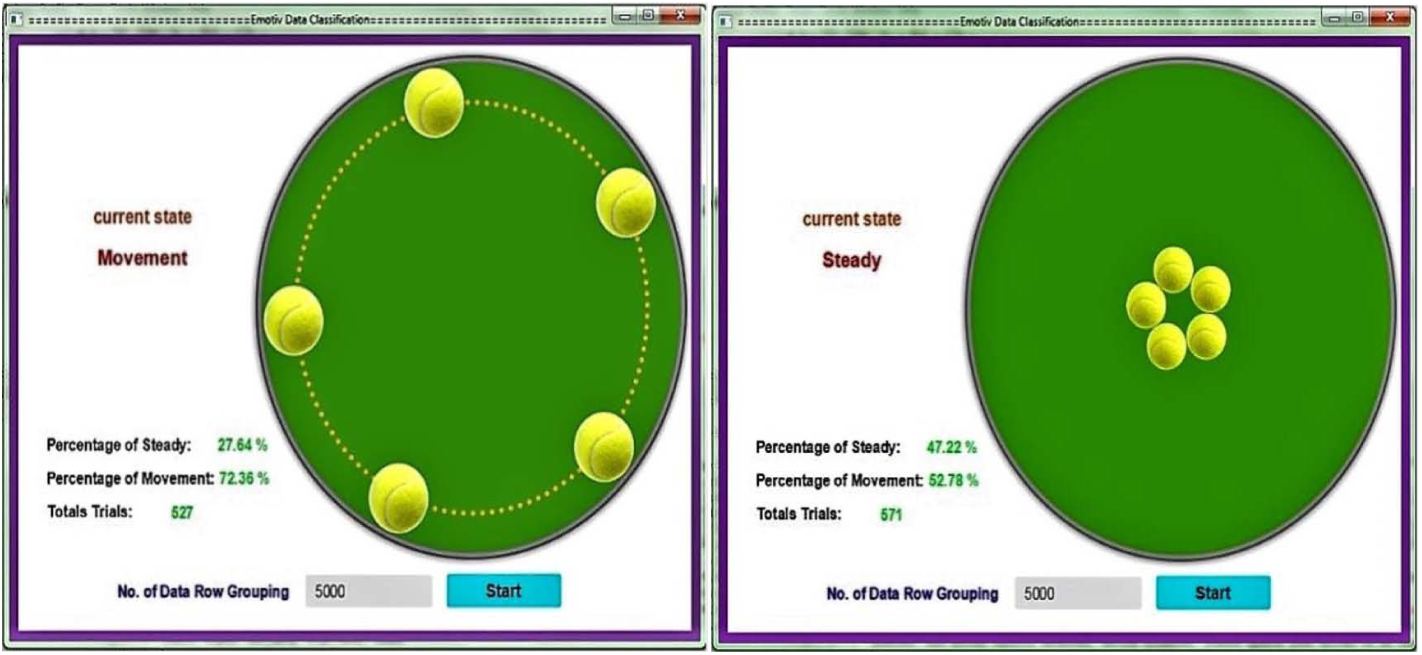
Classify real-time motor imagery tasks via developed brain game, presenting movements and steady correspondingly.

## 6. Conclusions & Future Work

In our research, we have used Emotiv Epoc+ EEG neuroheadset after analyzing several BMI devices like Emotiv Epoc+, Muse Headband, Aurora Dream, and MindWave. To obtain MI-EEG brain signals, we built an application program manipulating Emotiv SDK with Java technology. We have constructed several classifiers using ANN, SVM, naïve Bayes, Decision Tree, Random Forest, Bagging, AdaBoost and compared the performance of these existing approaches with the proposed clustering-based ensemble technique. The ensemble method we suggested, produced better performance than the above-mentioned classifiers. Then, the suggested method was used to develop the aimed application system. The game that we developed is capable of controlling the movements of the balls utilizing the real-time MI-EEG brain signals. It offers the user to enhance the quality of attention, which boosts productivity as well as upgrades the standard of works. It also assists people who are physically impaired or disabled and carries the potentials for human functionality enhancement. We designed the game to be a single-player game and to recognize three actions. In the future, more actions and more players can be added to make the game more advanced and challenging. The finding of this research can be applied to manipulate and enhance the control as well as movements of robots. It also brings new potentials in the health and rehabilitation industry.

